# Single cell transcriptional profiling reveals cellular diversity, communication, and sexual dimorphism in the mouse heart

**DOI:** 10.1101/201970

**Authors:** Daniel A. Skelly, Galen T. Squiers, Micheal A. McLellan, Mohan T. Bolisetty, Paul Robson, Nadia A. Rosenthal, Alexander R. Pinto

## Abstract

Characterization of the cardiac cellulome—the network of cells that form the heart—is essential for understanding cardiac development and normal organ function, and for formulating precise therapeutic strategies to combat heart disease. Recent studies have challenged assumptions about both the cellular composition^1^ and functional significance of the cardiac non-myocyte cell pool, with unexpected roles identified for resident fibroblasts^2^ and immune cell populations^3,4^. In this study, we characterized single-cell transcriptional profiles of the murine non-myocyte cardiac cellular landscape using single-cell RNA sequencing (scRNA-Seq). Detailed molecular analyses revealed the diversity of the cardiac cellulome and facilitated the development of novel techniques to isolate understudied cardiac cell populations such as mural cells and glia. Our analyses also revealed networks of intercellular communication as well as extensive sexual dimorphism in gene expression in the heart, most notably demonstrated by the upregulation of immune-sensing and pro-inflammatory genes in male cardiac macrophages. This study offers new insights into the structure and function of the mammalian cardiac cellulome and provides an important resource that will stimulate new studies in cardiac cell biology.

## MAIN TEXT

To acquire a high resolution map of the cardiac cellulome, we prepared a single-cell suspension of viable, metabolically active, nucleated non-myocyte cells from heart ventricles of female and male mice, for scRNA-Seq analysis. To minimize oversampling of endothelial cells, which form the largest cardiac cell population and principally comprise microvascular endothelial cells^1^, we reduced the proportion of endothelial cells to ~10% of total non-myocytes (Fig. 1A). We obtained transcriptional profiles of single cells using the 10X Chromium^TM^ platform, and analysed 10,519 cells passing quality control and filtering, for which an average of 1,897 genes per cell were measured (Fig. S1A).

**Figure 1.**
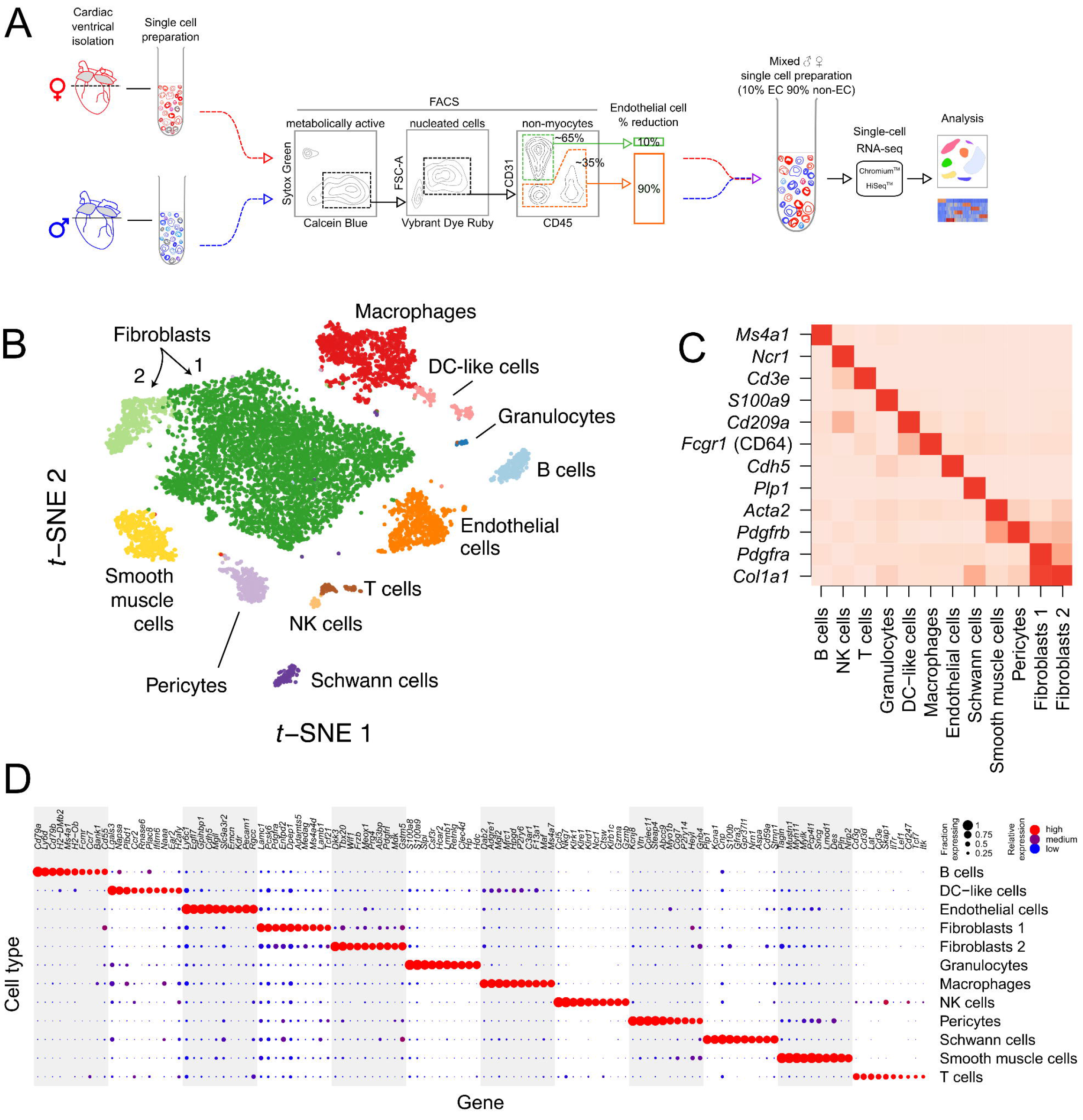
Single-cell RNA-Seq analysis of cardiac non-myocytes reveals major cardiac cell types. (A) Outline of experimental procedure for preparation of a non-myocyte, endothelial cell depleted cardiac cell preparation for single-cell RNA-Seq analysis. Fluorescence-activated cell sorting (FACS) was used to isolate viable (Sytox Green^-^), metabolically active (Calcein Blue^+^), nucleated (Vybrant DyeCycle Ruby^+^) cells and reduce the endothelial cell population as depicted (Materials and Methods). Single cell gating is shown in Fig. S3A. (B) Major cardiac cell populations identified following unsupervised clustering. Each point depicts a single cell, coloured according to cluster designation. We reduced dimensionality by PCA, used a graph-based approach to cluster cells, and performed spectral t-SNE to reduce the data to two dimensions for visualization (Materials and Methods). (C) Known cell type markers strongly and specifically associate with major cell types. Intensity of red hue in individual squares comprising the heatmap is proportional to the mean expression of each gene marked along the y-axis within all cells identified as the cell type noted on the x-axis. (D) Dot plot of ten marker genes for each major cell population. Marker genes were identified in an unbiased fashion blind to known cell type markers. Individual dots are sized to reflect the proportion of cells of each type expressing the marker gene, and coloured to reflect the mean expression of each marker gene across all cells, as indicated in the legend.

To identify distinct cell populations based on shared and unique patterns of gene expression, we performed dimensionality reduction and unsupervised cell clustering using methods implemented in the Seurat software suite (Materials and Methods)^5,6^. This clustering approach requires some user specification of parameters but is agnostic to known or predicted cell type markers. We identified twelve distinct cell clusters expressing known markers of major cell types (Fig. 1B-C). These comprised endothelial cells (*Cdh5, Pecam1*), fibroblasts (2 clusters; *Colal*, *Pdgfra*, *Tcf21*), granulocytes (*Ccr1*, *Csf3r*, *S100a9*), lymphocytes (3 clusters; *Ms4a1*, *Cd3e*, *Klrb1c*, *Ncr1*), pericytes (*P2ry14*, *Pdgfrb*), macrophages (*Adgre1*, *Fcgr1*), dendritic cell (DC)-like cells (*Cd209a*), Schwann cells (*Plp1*, *Cnp*), and smooth muscle cells (SMCs; *Acta2*, *Tagln*). Prototypical markers that define cell populations (Fig. 1C) included *Col1a1* (fibroblasts), *Ms4a1* (B lymphocytes), *Plp1* (Schwann cells) and *Acta2* (SMCs). We also identified additional genes that strongly and specifically marked each major cell population (Fig. 1D). Gene Ontology (GO) enrichment analysis of genes with restricted expression in each major cluster supported predicted identities of cell populations (Supplementary data file, Tab 1). For example, fibroblast distinctness is driven by extracellular modelling terms; leukocytes are associated with immune-regulatory terms; endothelial cells and SMCs are associated with vascular development and homeostasis terms; and Schwann cells are associated with neuro-regulatory terms.

To reveal heterogeneity within major cell types, we carried out subclustering of these populations (Fig. 2A; Fig. S1B-C). We used a similar unsupervised approach as described above, and analysed cells of a single type or multiple related types (e.g. myeloid or lymphoid leukocytes) to discover cell subtypes (Materials and Methods). For some cell populations, such as endothelial cells, Schwann cells, and smooth muscle cells, subclustering did not identify transcriptionally distinct subpopulations (Fig. 2B), suggesting that these populations are relatively homogeneous in the mature heart. In contrast, multiple levels of subpopulation heterogeneity emerged within other cell populations, where subclustering clearly and distinctly separated identifiable populations. For example, subclustering of lymphoid cell populations revealed a population of group 2 innate lymphoid cells (ILC2s)^7^, showing expression of genes such as *Gata3*, *Areg*, and *Rora*, that previously clustered with T lymphocytes (Fig. 1B; Fig. 2A-B). Moreover, further subclustering of T lymphocytes alone produced two clusters corresponding to CD4^+^ and CD8^+^ T cells (Fig 2A, inset).

**Figure 2.**
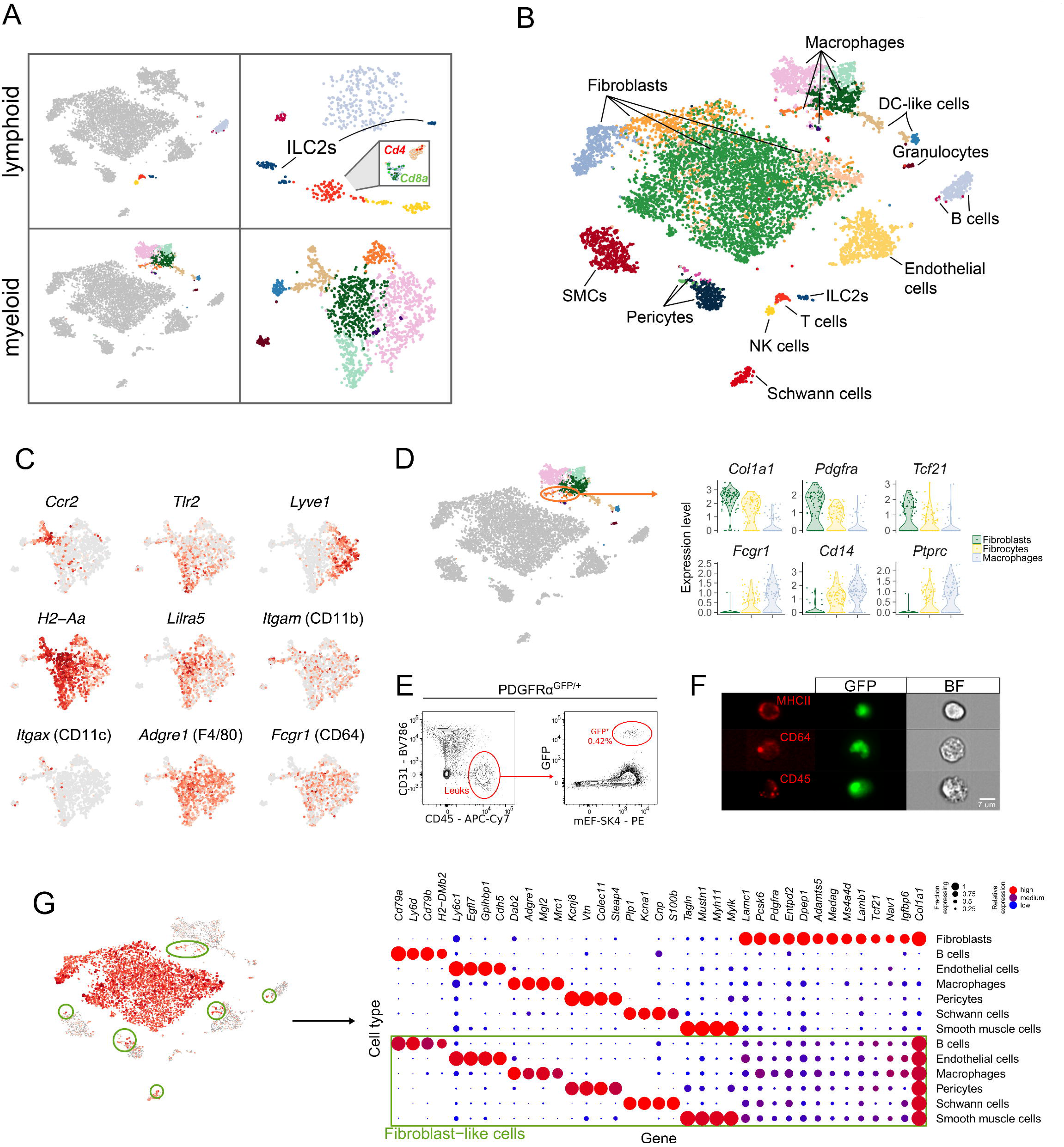
Major cardiac cell types harbour subpopulations that reflect a hierarchy of transcriptional diversity. (A) Subclustering of lymphoid (top) and myeloid (bottom) cell populations reveals structure that is not apparent when clustering at a global level (Fig. 1). Left panels depict cells positioned in t-SNE space with the populations of interest coloured and all other cells grey. Right panels are coloured to reflect subcluster designations and show cells positioned in subcluster t-SNE space. Inset shows tertiary clustering of T cells to identify CD4^+^ and CD8^+^ T cell populations. (B) Detailed map of cardiac cellular diversity following subclustering. Cells are coloured according to subcluster designation. (C) Gene expression gradients identified within macrophage/DC-like populations. Points show myeloid lineage cells positioned in t-SNE space identically to part (A), lower right, with more intense red hue indicating higher relative expression and grey signifying no expression. (D) Identification of putative fibrocytes that exhibit gene expression signatures of both fibroblasts and macrophages. Violin plots display gene expression of canonical fibroblast (*Col1a1*, *Pdgfra*, *Tcf21*) and macrophage/leukocyte (*Fcgr1*, *Cd14*, *Ptprc*) genes. (E) Identification of fibrocytes within the uninjured hearts of PDGFRα^GFP/+^ mice, where GFP expression principally identifies fibroblasts. (F) ImageStream^®^ cytometry analysis of GFP^+^ cells from PDGFRα^GFP/+^ mouse hearts labelling for MHCII (*H2-Aa/Ab*), CD64 (*Fcgr1*) and CD45 (*Ptprc*). (G) Major cell populations contain minor subsets of cells expressing fibroblast-like gene signatures. Left panel: expression of *Col1a1* (grey=low, red=high) in cells plotted in t-SNE space identically to panel (B), except diameter of each point is also proportional to expression level of *Col1a1*. Minor subsets of B cells, endothelial cells, macrophages, pericytes, Schwann cells, and smooth muscle cells expressing *Col1a1* are circled. Right panel: dot plot shows expression of a selection of cell-type specific genes in subsets of cardiac cells. Data outlined in green highlights fibroblast-like cells in these groups by gating *in silico* on *Col1a1* expression. Each row is based on a random sample of 35 cells from the cell type shown. Individual dot size and colour reflects the proportion of each cell type expressing marker genes, and mean expression of the gene across all cells (indicated by legend).

In some major clusters, transcriptional heterogeneity that differentiated subpopulations reflected continuous gradients rather than discrete groups of homogeneous cells with clear gene expression boundaries. For example, subclustering all myeloid populations together identified the granulocyte population detected at the global level (Fig. 1B-C), but macrophage subtypes were less clearly separable and were connected by transcriptional gradients (Fig. 2A-C; Fig. S2). This presents a more nuanced picture of the macrophage transcriptional landscape than offered by the canonical subtyping of cardiac tissue macrophages based on markers such as *Cx3cr1*, *H2-Aa/Ab* (MHCII), *Ccr2*, *Mrc1* and *Adgre1* (F4/80)^8–10^. At least five subtypes of macrophages can be classified in our data on the basis of subtle differences in the expression of combinations of several genes (Fig. 2A-C), and in some cases additional myeloid subpopulations can be discovered following further subclustering (e.g. DC cell-like subtypes; Fig. S2). Similar transcriptional continua have also been identified in other tissues using scRNA-Seq^11^. This blending of profiles to form intermediate populations could represent transcriptional heterogeneity that is buffered at the protein level. Alternatively, these hybrid expression signatures could reflect the multifaceted nature of macrophages, with cells poised to respond to a diverse range of stimuli.

Among macrophages, we discovered a subpopulation of cells that exhibit a hybrid molecular signature of both macrophages and fibroblasts, resembling a putative fibrocyte population^12,13^. These cells express intermediate levels of canonical genes corresponding to both fibroblasts (*Col1a1*, *Pdgfra*, *Tcf21*) and macrophages/leukocytes (*Fcgr1*, *Cd14* and *Ptprc*) (Fig. 2D). This observation is not due to technical artefacts such as cell doublets or ambient RNA contamination: the population's size is larger than would be expected based on doublet rate estimates (<5%; 10X Genomics), and ambient RNA contamination would likely give rise to transcriptional profiles overlapping at least one of the most abundant cardiac cell types, namely cardiomyocytes or endothelial cells (Pinto et al. 2016). We confirmed the presence of putative fibrocytes by detecting GFP^+^ leukocytes in PDGFRα^GFP/+^ mouse hearts (Fig. 2E), and using ImageStream^®^ cytometry to capture images of GFP^+^ single cells bearing macrophage surface markers (Fig. 2F). It is noteworthy that we also detected minor cell subsets exhibiting fibroblast-like gene signatures within several other major cell populations^14^ (Fig. 2G) that appeared genuine. For example, we observed *Icam2*^+^ endothelial cells showing fibroblast-like gene expression, which we confirmed with cytometric images of GFP^+^ single cells expressing ICAM2 (Fig. S3E). Therefore, in line with well-established mesenchymal transition paradigms^15^, expression of a hybrid mesenchymal phenotype is likely an inherent characteristic of a broad range of cell types.

Amongst the major cardiac cell populations identified in the present analysis, notable absences of known cardiac cell populations included epicardial cells marked by *Tbx18*, expression of which was dispersed amongst cells falling within fibroblast and mural cell clusters (data not shown). *Isl1* transcripts marking putative cardiac progenitors were present in four cells, but these were dispersed across various cardiac cell populations and did not form a distinct cluster (data not shown). Lymphatic endothelial cells expressing *Prox1* and *Pdpn* were also absent from the artificially reduced endothelial population, but these cells normally comprise only ~3 % of total non-myocytes^1^. We also did not detect a discrete population of monocytes, which are present at low frequency and are likely dispersed among macrophage clusters. Overall, the absence of these cell populations is likely due to their relatively low abundance and lack of highly distinctive transcriptional signatures.

Mural cells (pericytes and SMCs) are essential components of vasculature^16,17^, but tools have been lacking for precise isolation of these cells by flow cytometry. Similarly, although Schwann cells are known to be present in the heart^18^, flow cytometric approaches to isolate these cells are limited. Commonly used markers including *Cspg4* (also called NG2), *Pdgfrb* and *Mcam* are relatively non-specific (Fig 3A); *Cspg4* and *Mcam* are expressed in mural cells and Schwann cells, and *Pdgfrb* expression is detected in mural cells and fibroblasts (Fig 3A). To develop strategies to discriminate pericytes, SMCs, Schwann cells, and fibroblasts, we identified genes that showed higher expression in one of these cell types relative to the others using a likelihood ratio-based test^19^ implemented in Seurat^5^. Genes identified using this approach with commercially available antibody reagents included *Itga7*, *Entpd1* and *Cd59a* (Fig. 3A; Supplementary data file, Tab 2). Examination of resident mesenchymal cells (RMCs; Fig. S3A) following ITGA7 staining enabled mural cells to be distinguished from fibroblasts using mEF-SK4^1^ as a secondary marker. This approach identified three populations: R1 (mEF-SK4^hi^ITGA7^-^); R2 (mEF-SK4^int/lo^ITGA7^+^); and R3 (mEF-SK4^lo^ITGA7^-^) (Fig 3C). Analysis of PDGFRα^GFP^ and CSPG4-GFP mice that mark expression in fibroblasts^20^ and mural cells^21^ respectively (Fig 3C-D), revealed that almost all GFP^+^ resident mesenchymal cells (RMCs) in the PDGFRα^GFP/+^ mouse hearts are within R1, whereas almost all GFP^+^ RMCs in CSPG4-GFP mouse hearts are within R2 (Fig. 3D). Conversely, analysis of GFP^+^ cells in R1-3 reveals that almost all R1 cells are GFP^+^ in PDGFRα^GFP^ mouse hearts, whereas the majority of R2 cells are GFP^+^ in CSPG4-GFP mice (Fig. 3C). Thus, cardiac fibroblasts (defined here as PDGFRα^+^ cells) can be identified and isolated with high precision using the mEF-SK4 and ITGA7 markers, where ITGA7 distinguishes mural cells from mEF-SK4^low/int^ fibroblasts. Similar to ITGA7, we found that MCAM separates mural cells from fibroblasts, but also marks Schwann cells. In addition, we validated that ENTPD1 and CD59a identify SMCs and Schwann cells, respectively (Fig. S3C-D). By combining these markers, we were able to identify and isolate SMCs, pericytes, and Schwann cells (Fig. 3F-G).

**Figure 3.**
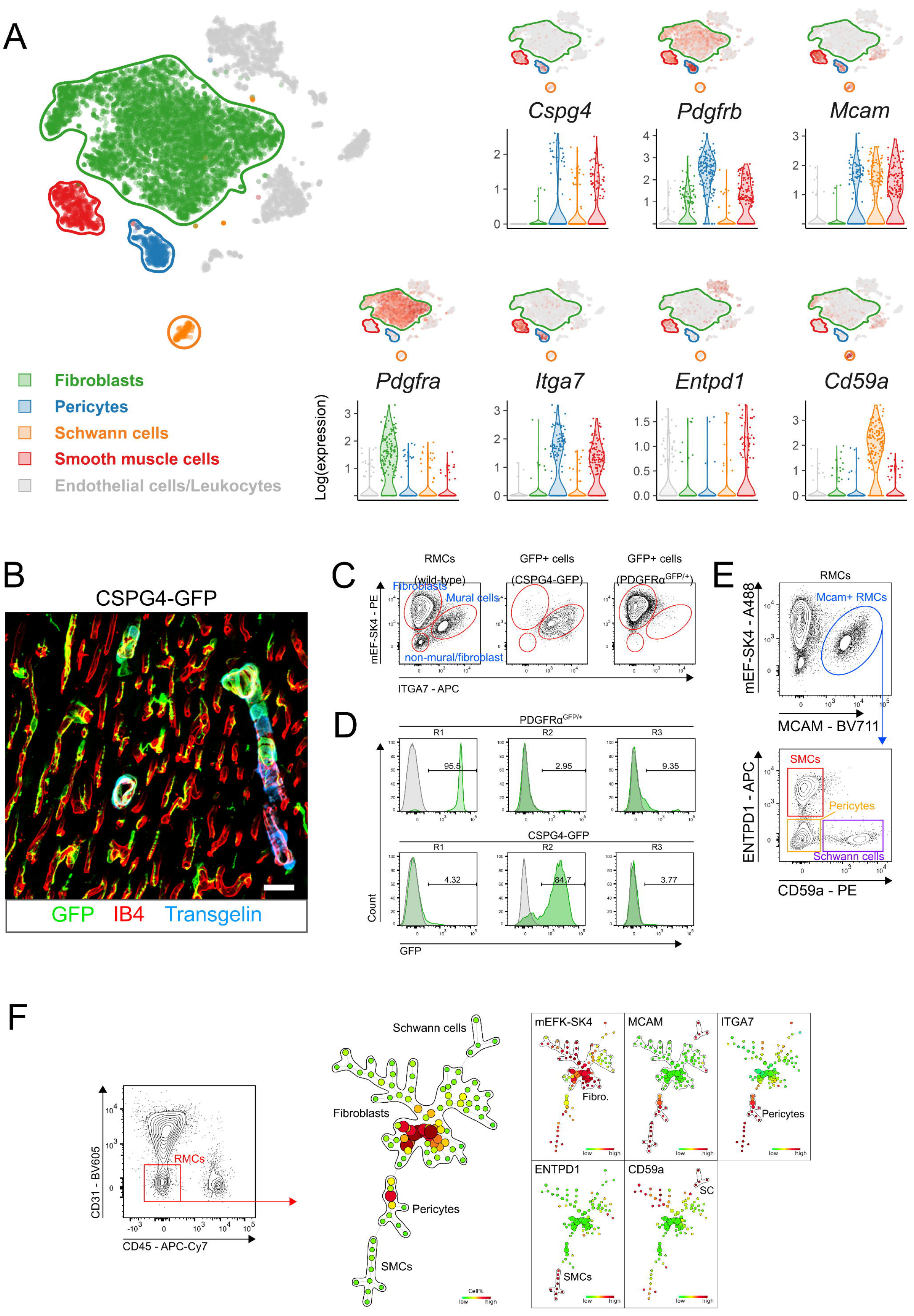
Novel strategies for isolation of cardiac mural cells and glia. (A) Plots showing cell position in t-SNE space along with expression of select genes expressed in cardiac mural cells and glia. Genes shown include both existing and novel markers to identify resident mesenchymal cell (RMC, non-endothelial CD31^+^, non-leukocyte CD45^+^ cells; Fig. S3A) subpopulations including mural cells (pericytes and SMCs). Upper left plot shows clustering as in Fig. 1 but with only fibroblast, glia, and mural cell populations highlighted. Smaller t-SNE plots show expression of select genes with populations outlined as to the left but with cells coloured according to the relative expression of the gene shown (red=high, grey=low). Violin plots below t-SNE plots were constructed using data obtained from 150 randomly sampled cells from each major population and show expression of each gene broken down by cell population as indicated in the legend at lower left. (B) Projection view of GFP expression in CSPG4-GFP mouse hearts. Sections are also stained for transgelin and isolectin B4 (IB4) to show both pericytes (GFP+transgelin^-^) and SMCs (GFP^+^transgelin^+^). Scale bar indicates 20μm. (C) Discrimination of fibroblasts and mural cells using anti-ITGA7 and anti-mEF-SK4 antibodies. Left panel, RMCs were pre-gated as shown in Fig. S3A. Middle and right panels, cells were pre-gated GFP^+^ RMCs. Text in parenthesis indicates genotypes of mice from which heart cells were prepared. (D) Histograms of GFP^+^ cells within fibroblast, mural cell and non-fibroblast/mural cell populations shown in (C). Text above plots indicates genotypes of mice from which heart cells were prepared. (E) Identification of SMCs, pericytes and Schwann cells using MCAM, CD59a and ENTPD1. (F) SPADE dendrogram demonstrating that fibroblasts, mural cells, and Schwann cells show distinct signatures that allow for their precise isolation. Each circle (‘node’) represents a phenotypically similar/identical cell population based on surface markers (shown on right) used for clustering. Colour and size of circles are proportional to abundance of cell population, and node connections indicate phenotypic similarity of neighbouring nodes.

To define intercellular communication networks within the cardiac cellulome we utilized a dataset of human ligand-receptor pairs^22^ to develop a list of mouse orthologues comprising 2,009 ligand-receptor pairs (Supplementary data file, Tab 3; Materials and Methods). Although anatomical barriers between cell types are not modelled in this analysis, expression patterns of ligand-receptor pairs in each cell type revealed a dense intercellular communication network (Fig 4A). GO enrichment analysis of ligands for which cognate receptors were present in cardiac non-myocytes revealed genes involved in cell positioning (locomotion, migration, etc.), expressed by non-patrolling cardiac cell populations (non-B, -T cells, or granulocytes), while analysis of receptors showed enrichment for processes such as cell communication and signal transduction (Supplementary data file, Tab 4). Broadcast ligands for which cognate receptors are detected within cardiac non-myocytes (Fig. 4A) identified fibroblasts as the most trophic cell population with dense connections to multiple cell types (Fig. 4B). These include signalling circuits that support survival of specific cardiac cell populations (Fig. 4C). For example, fibroblasts and pericytes express *Csf1* and *Il34*, respectively (Fig. 4D), which signal through CSF1R and are essential factors for macrophage growth and survival. Fibroblasts also express growth factors *Ngf*, *Ntf3*, *Vegfa*, *Igf1* and *Fgf2* (Fig. 4C) which support neurons of the autonomous nervous system, endothelial cells, and mural cells^23–26^

**Figure 4.**
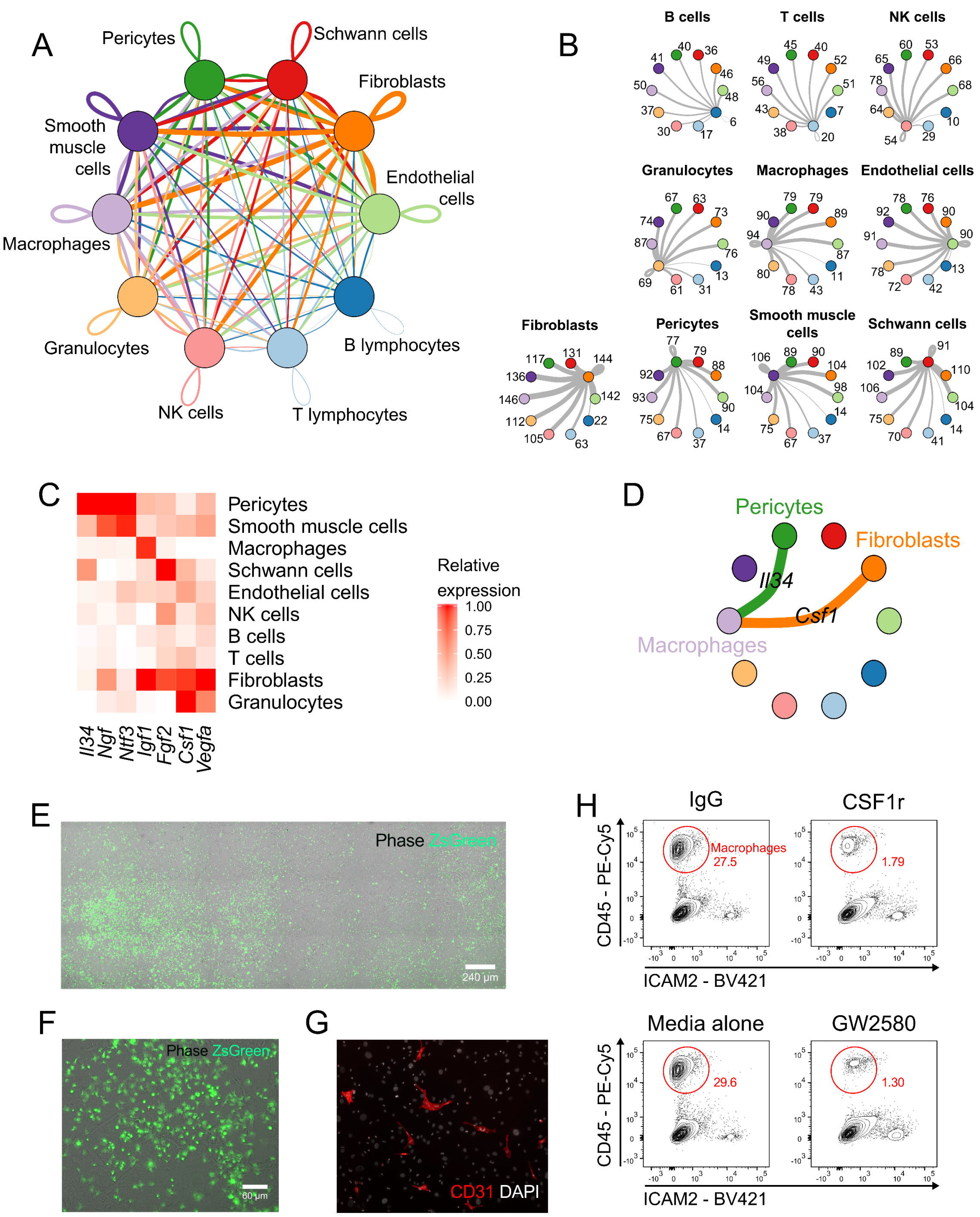
A dense network of autocrine and paracrine signalling underpins cardiac homeostasis. Two fibroblast populations, as well as DC-like cells and macrophages, have ligand and receptor profiles that are substantially similar and are merged for clarity. (A) Capacity for intercellular communication between cardiac cell types. Line colour indicates ligands broadcast by the cell population of the same colour (labelled). Lines connect to cell populations where cognate receptors are expressed. Line thickness is proportional to the number of ligands where cognate receptors are present in the recipient cell population. Loops indicate autocrine circuits. Map quantifies potential communication but does not account for anatomic position or boundaries of cell populations. (B) Detailed view of ligands broadcast by each major cell population and those populations expressing cognate receptors primed to receive a signal. Numbers indicate quantity of ligand-receptor pairs for each inter-population link. (C) Production of essential growth factors by cardiac cell populations. (D) Non-patrolling cardiac cell populations supporting macrophage growth by CSF1 or IL34 production. (E) Composite micrograph displaying phase contrast image of mixed cardiac non-myocyte cell culture and ZsGreen fluorescence (green). Non-myocyte cells were prepared from a LysM-Cre:Rosa^Zsgreen/+^ hearts where myeloid leukocytes are labelled with ZsGreen. Images taken two weeks post-plating. (F) Magnified view of cells shown in (E). (G) Fluorescence micrograph of CD31^+^ endothelial cells (red) and DAPI^+^ nuclei (white) in mixed non-myocyte cultures, two weeks post-plating. (H) Macrophage growth *in vitro* is obstructed by inhibition of CSF1R signalling. Flow cytometry contour plots show the effect of inhibition of CSF1R signalling by anti-CSF1R antibody or using synthetic compound GW2580. Flow cytometric gating for generation of contour plots is shown in Fig. S3B.

To examine the capacity of cardiac cells to support cell types within the cardiac cellulome, we cultured mixed cardiac cells and observed growth of disparate cell populations *in vitro*. Non-myocyte cultures from LysM-Cre:Rosa^ZsGreen/+ 27,28^ mice generated abundant macrophages labelled by ZsGreen fluorescent protein (Fig. 4E-F), as well as endothelial cells (Fig. 4G). However macrophages failed to grow when cultures were treated with a CSF1R-blocking antibody^29^, synthetic CSF1R inhibitor GW2580^30^ (Fig. 4H), or when cultured in normal growth media without non-macrophage cells (data not shown). Although these results indicate that fibroblasts support both cardiac macrophage and endothelial cell growth, non-fibroblast sources of essential growth factors point to the complexity of the network of cells that establish the cardiac niche and support resident cell populations.

Our analyses also show that diverse cell populations support nervous innervation of the heart. Consistent with recent findings that cardiac nerves develop along vasculature^31^, we found that mural cells express *Ngf* and *Ntf3*, both important factors for axonal development (Fig. 4C). Expression of *Ngf* and *Ntf3* by cardiac pericytes highlights their potentially important role in development of the autonomic nervous system in the heart. Moreover, cardiac Schwann cells, the myelin-producing cells of the peripheral nervous system, are a discrete cardiac cell population^18^. Thus, multiple cardiac cell types contribute to supporting the development, maintenance, and signal transduction of autonomic nerves.

To examine sexual dimorphism in cardiac non-myocyte gene expression, we segregated female and male cells based on expression of female- (*Xist*) and male-specific genes (six Y chromosome genes: *Ddx3y*, *Eif2s3y*, *Erdr1*, *Gm29650*, *Kdm5d*, *Uty*; Fig 5A). Highly similar clustering patterns for female and male cells were observed at the global level, but within individual cell types we identified many genes showing sexual dimorphism in their expression (Fig. 5A). Of the 396 genes upregulated in female cells and 430 upregulated in males (Materials and Methods), the vast majority (~97%) were differentially expressed below 2-fold (Fig. 5B), with a median fold difference of 1.17 for both females and for males. The mechanisms underlying sexual dimorphism are likely to be diverse and encompass both subtle shifts in composition of cell subtypes as well as regulatory responses to hormonal cues or trans-acting sex chromosome factors.

**Figure 5.**
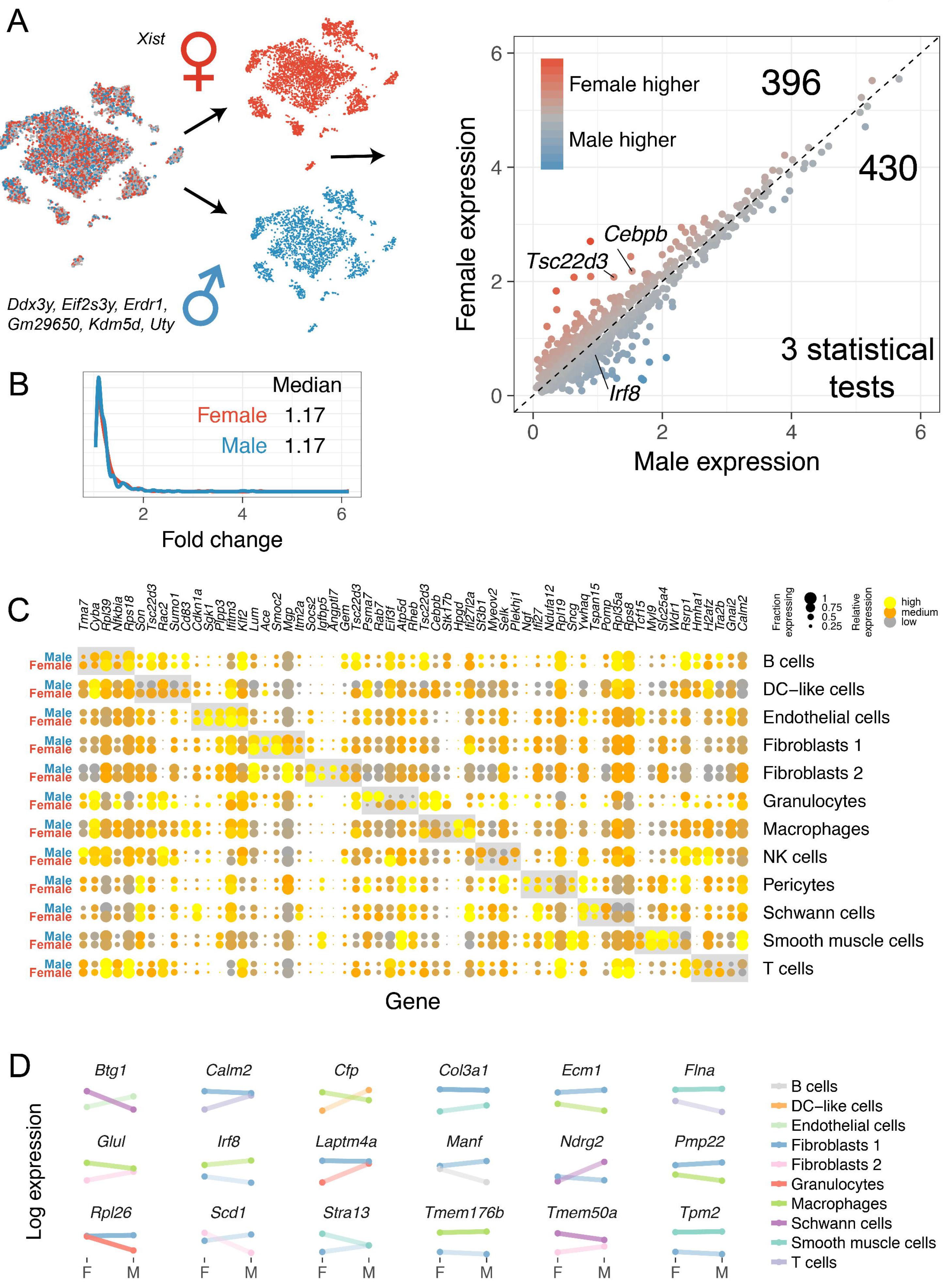
Widespread cell type-specific sexual dimorphism in cardiac gene expression. (A) Left panel: identification of male and female cells using expression of *Xist* and six Y chromosome genes. Right panel: scatter plot of sexually dimorphic genes identified within each cell type (p < 0.05, three statistical tests; see Methods). Sex-linked genes are not shown. Numbers at upper right indicate count of genes upregulated in specific cell types in females and in males. Units on *x*- and y-axis are UMIs per ten thousand. (B) Histogram of fold change estimates for sexually dimorphic genes that are more highly expressed in females (red) and males (blue). (C) Dot plot showing top sexually dimorphic genes (up to five for each major cell population; grey boxes) and expression of the same genes stratified by sex in all cell populations. (D) Drumstick plot showing sexually dimorphic genes with discordant directionality in different cell populations. Points indicate mean expression of female and male cells according to colours in legend.

Analysis of the most highly sexually dimorphic genes for each cell population provided clear evidence that the extent and direction of sexual dimorphism in gene expression is cell type dependent (Fig. 5C; Supplementary data file, Tab 5). Furthermore, we identified 27 genes that exhibited discordant patterns of sexually dimorphic expression between cell types (Fig. 5D), revealing the dichotomous effect of biological sex on gene expression in the heart. Among male macrophages, we found evidence that male-upregulated genes tend to play a role in responding to foreign antigens, with significant GO enrichment for terms including antigen processing and presentation via MHC class II molecules as well as the broader immune response (adjusted *p* < 0.001; Supplementary data file, Tab 6). In contrast, female-upregulated genes in macrophages were enriched for processes involving response to stress and the electron transport chain (Supplementary data file, Tab 6).

These results support epidemiological and experimental data documenting sexually dimorphic responses to cardiac insults^32^. Females exhibit cardioprotection with decreased neutrophil infiltration and reduced cardiomyocyte death^33,34^, whereas the present analysis reveals that the male cardiac cellulome is particularly geared towards sensing inflammatory cues. We observe sexual dimorphism in cardiac tissue macrophage transcriptional profiles, such as upregulation of *Irf8*, which is linked to chronic inflammation^35^. Conversely, the most upregulated gene in female macrophages, and the most sexually dimorphic macrophage gene in both sexes (Fig. 5C), is *Tsc22d3* (also known as Gilz), a transcription factor implicated in anti-inflammatory functions and a downstream driver of the potent anti-inflammatory effects of glucocorticoids^36–38^. The functional significance of this gene has been demonstrated by the transplantation of cells overexpressing *Tsc22d3* to infarcted murine hearts which results in dampened inflammation and reduced cardiomyocyte cell death^39^. However, it is unlikely that macrophage *Tsc22d3* expression alone confers the anti-inflammatory phenotype in female hearts, which also have elevated expression of other genes implicated in anti-inflammatory mechanisms, such as *Cebpb*^40^ (Fig. 5C).

This study builds on previous explorations of cardiac cellular diversity^1,14^ and provides unique insights into the structure and function of the cardiac cellulome. We have characterized single-cell transcriptional profiles of major non-myocyte cell populations within the mouse heart. We observed a dense ligand-receptor network facilitating extensive communication between diverse cell types and documented a significant contribution of biological sex to the transcriptional programs that govern organ function. An important aspect of the dataset presented here is its utility for identifying and studying cardiac cells. Here we used our scRNA-Seq dataset to develop strategies to more precisely identify cardiac mural cells, fibroblasts, and Schwann cells using flow cytometry. Our results will stimulate research into new avenues of inquiry in cardiac cell biology and provide a resource for in-depth interrogation of the cell type-specific transcriptional networks that underpin heart development, homeostasis, and disease.

## ACKNOWLEDGEMENTS

A.R.P. is supported by an American Heart Foundation Grant (17IRG33270004). The Australian Regenerative Medicine Institute is supported by grants from the State Government of Victoria and the Australian Government. The authors are grateful for assistance with library preparation and sequencing from the Jackson Laboratory Genome Technologies facility.

## AUTHOR CONTRIBUTIONS

D.A.S and A.R.P wrote the manuscript with input from all co-authors. A.R.P., with technical assistance from P.R. and D.A.S., designed experiments. M.T.B., with supervision by P.R., analysed raw data. G.T.S., M.A.M., and A.R.P. carried out cell preparation for RNA-Seq and conducted flow cytometry experiments. M.A.M. and A.R.P carried out cell culture experiments. A.R.P., G.T.S., and M.A.M analysed flow cytometry data. D.A.S. analysed all scRNA-Seq data. N.A.R. and A.R.P. provided supervision and guided presentation of the data. A.R.P conceived the project.

## SUPPLEMENTARY FIGURE LEGENDS

**Figure S1.**
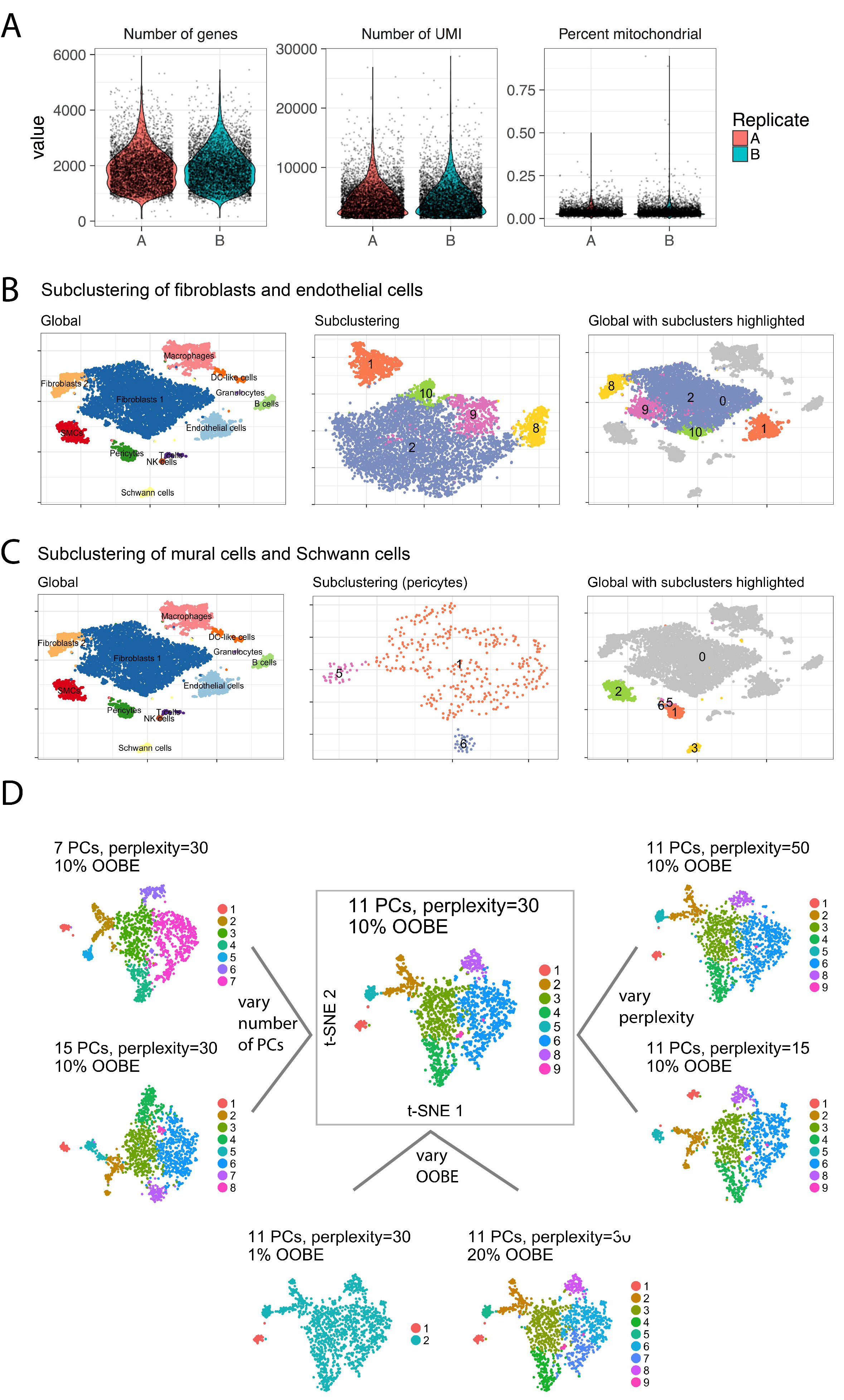
Quality control, subclustering, and clustering parameter choices for scRNA-Seq data. (A) Violin plots showing number of genes, UMIs, and percent mitochondrial reads for all cells in the full dataset. (B) Subclustering results for fibroblast and endothelial cells. Cells within these populations were separately isolated from the full dataset and reclustered (Materials and Methods). Left panel recapitulates primary clusters (Fig. 1B). Middle panel indicates positioning of cells in subcluster t-SNE space. Right panel shows cells in same layout as at left, but with cells coloured according to subcluster designation. (C) Subclustering results for SMCs, pericytes, and Schwann cells. Plots are the same as in (B) but in the middle panel only pericytes are shown for clarity as they were the only population that separated during subclustering. (D) Robustness of clustering results to variation in parameter choices. As an illustration, we show the effect of varying parameter choices on subclustering of myeloid lineage cells. Each t-SNE plot shows the effect of varying parameters as shown in the title above the plot. Central plot represents the parameters chosen for myeloid subclustering presented in the main text.

**Figure S2.**
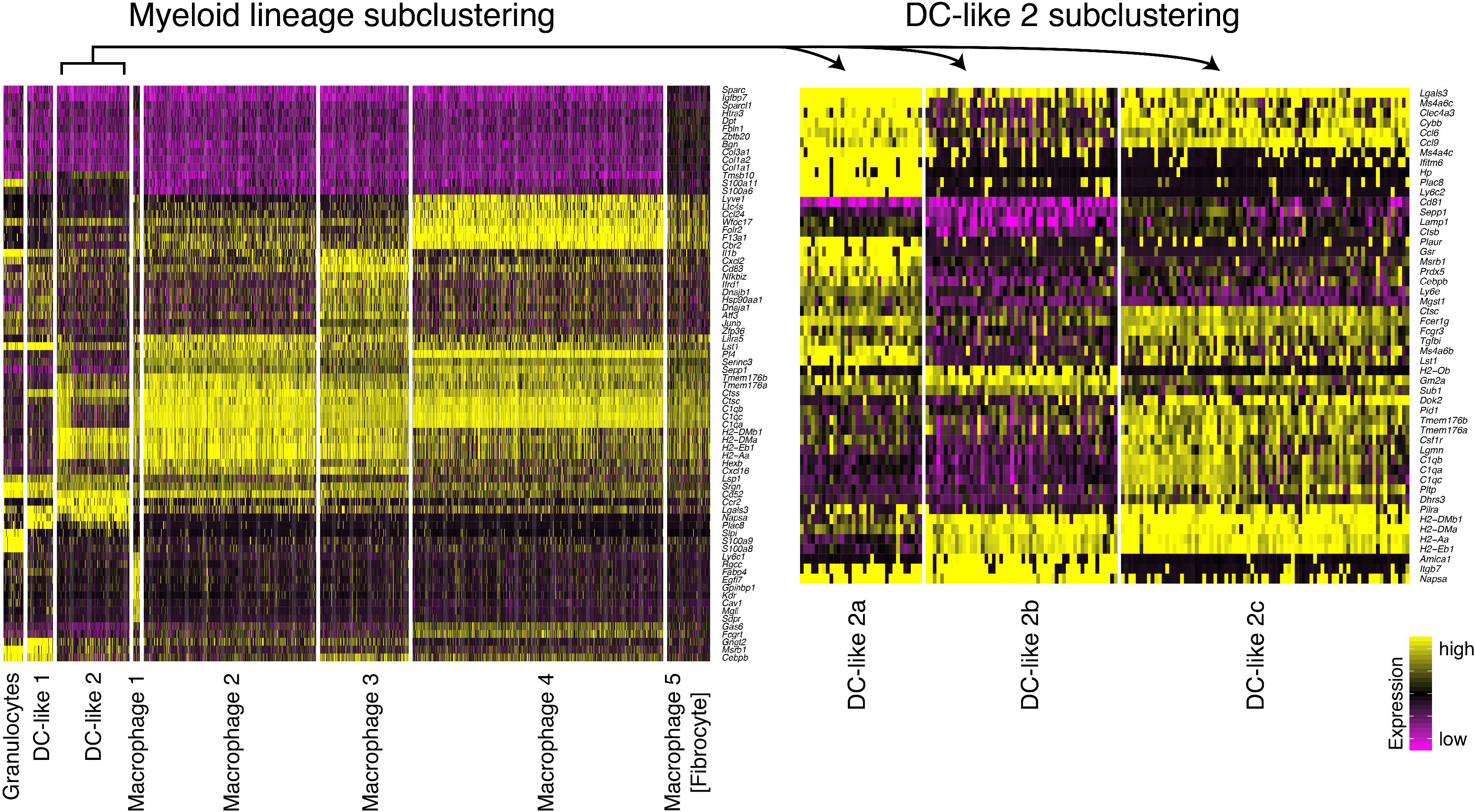
Subclustering reveals transcriptionally distinct subsets of the myeloid lineage. Left panel shows heatmap of genes (rows) that best distinguish between eight myeloid lineage subclusters (columns). The DC-like 2 subcluster shows notable gradients across several genes. Upon isolating this cell subset and re-clustering, we identified an additional three subclusters (right panel). Colouring of gene expression is relative and is based upon expression of the gene in all cells in the full dataset.

**Figure S3.**
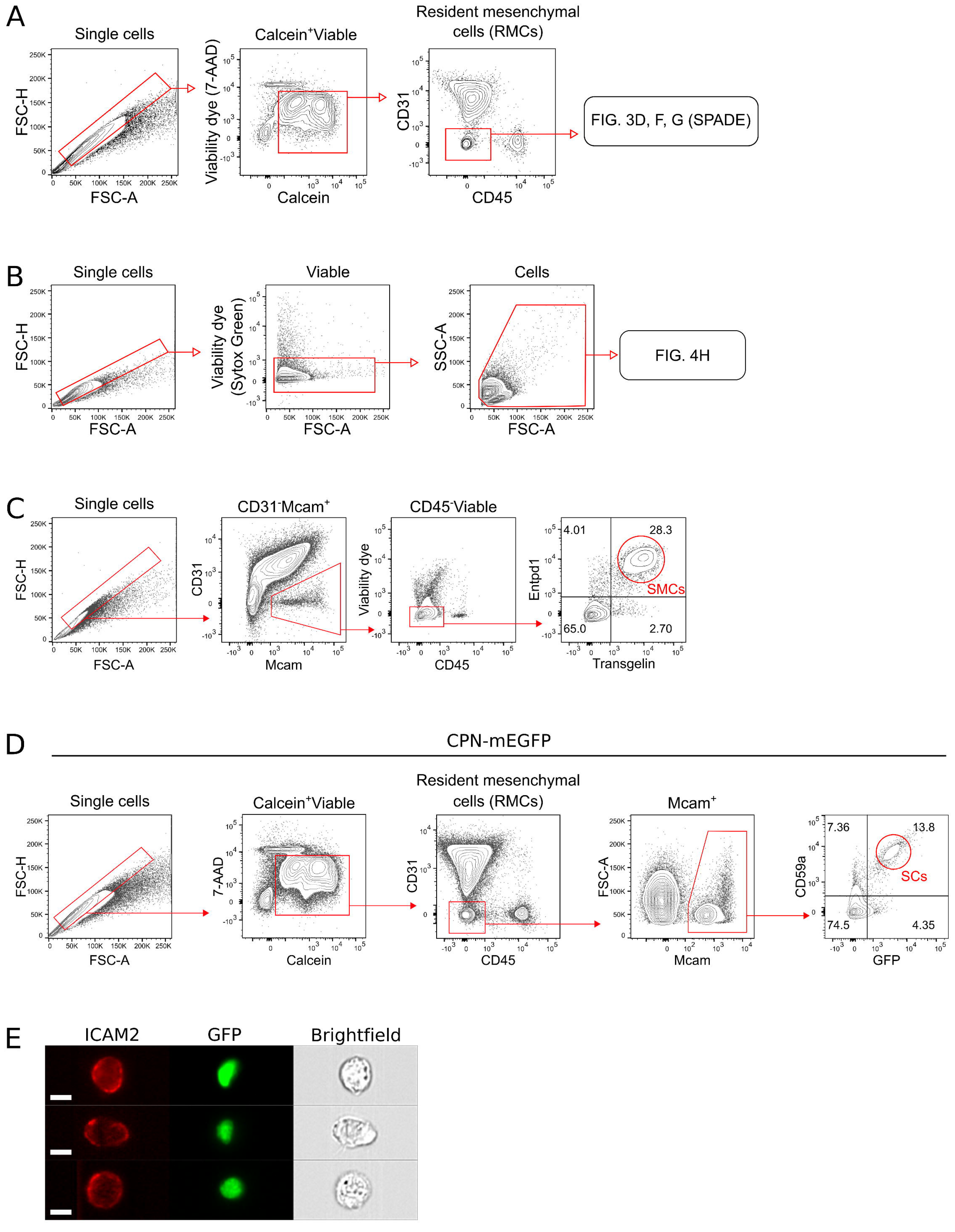
Flow cytometry gating and marker validation. (A) Gating strategy of viable (Calcein^+^7-AAD^-^) single cells preceding gating of resident mesenchymal cells (RMCs; CD31^-^CD45^-^ cells). (B) Gating strategy for identification of viable (Sytox Green^-^), single cells used for identifying cultured cell populations. (7-AAD: 7-aminoactinomycin D; FSC-H: forward scatter height; FSC-A: forward scatter area; SPADE: spanning-tree progression analysis of density-normalized events). (C) Validation of ENTPD1 labelling of smooth muscle cells. Contour plots show gating strategy and overlapping ENTPD1 staining with the smooth muscle cell marker transgelin following fixation and intracellular staining. (D) Validation of CD59a labelling of cardiac Schwann cells. Contour plots show gating strategy and overlapping labelling of CD59a staining with GFP in CPN-mEGFP mice. CPN-mEGFP mice express a membrane-localizing green fluorescent protein in oligodendrocytes and Schwann cells. (E) ImageStream® cytometry micrographs showing GFP^+^ cells from PDGFRα^GFP/+^ mouse hearts that express the endothelial cell marker ICAM2. Scale bar indicates 7 μm.

## REFERENCES

1. Pinto, A. R. et al. Revisiting cardiac cellular composition. Circ. Res. 118, 400–409 (2016).

2. Furtado, M. B. et al. Cardiogenic genes expressed in cardiac fibroblasts contribute to heart development and repair. Circ. Res. 114, 1422–1434 (2014).

3. Hulsmans, M. et al. Macrophages Facilitate Electrical Conduction in the Heart. Cell 169, 510–522.e20 (2017).

4. Monnerat, G. et al. Macrophage-dependent IL-1β production induces cardiac arrhythmias in diabetic mice. Nat. Commun. 7, 13344 (2016).

5. Macosko, E. Z. et al. Highly parallel genome-wide expression profiling of individual cells using nanoliter droplets. Cell 161, 1202–1214 (2015).

6. Butler, A. & Satija, R. Integrated analysis of single cell transcriptomic data across conditions, technologies, and species. bioRxiv (2017). doi:10.1101/164889

7. Artis, D. & Spits, H. The biology of innate lymphoid cells. Nature 517, 293–301 (2015).

8. Epelman, S. et al. Embryonic and Adult-Derived Resident Cardiac Macrophages Are Maintained through Distinct Mechanisms at Steady State and during Inflammation. Immunity 40, 91–104 (2014).

9. Pinto, A. R. et al. An abundant tissue macrophage population in the adult murine heart with a distinct alternatively-activated macrophage profile. PLoS One 7, e36814 (2012).

10. Ilinykh, A. & Pinto, A. R. The role of cardiac tissue macrophages in homeostasis and disease. Advances in Experimental Medicine and Biology 1003, (Springer, Cham, 2017).

11. Gokce, O. et al. Cellular Taxonomy of the Mouse Striatum as Revealed by Single-Cell RNA-Seq. Cell Rep. 16, 1126–1137 (2016).

12. Reilkoff, R. A., Bucala, R. & Herzog, E. L. Fibrocytes: emerging effector cells in chronic inflammation. Nat. Rev. Immunol. 11, 427–435 (2011).

13. Pilling, D., Fan, T., Huang, D., Kaul, B. & Gomer, R. H. Identification of markers that distinguish monocyte-derived fibrocytes from monocytes, macrophages, and fibroblasts. PLoS One 4, e7475 (2009).

14. DeLaughter, D. M. et al. Single-Cell Resolution of Temporal Gene Expression during Heart Development. Dev. Cell 39, 480–490 (2016).

15. Kovacic, J. C., Mercader, N., Torres, M., Boehm, M. & Fuster, V. Epithelial-to-mesenchymal and endothelial-to-mesenchymal transition from cardiovascular development to disease. Circulation 125, 1795–1808 (2012).

16. Gaéngel, K., Genové, G., Armulik, A. & Betsholtz, C. Endothelial-mural cell signaling in vascular development and angiogenesis. Arteriosclerosis, Thrombosis, and Vascular Biology 29, 630–638 (2009).

17. Armulik, A., Genové, G. & Betsholtz, C. Pericytes: Developmental, Physiological, and Pathological Perspectives, Problems, and Promises. Dev. Cell 21, 193–215 (2011).

18. Fregoso, S. P. P. & Hoover, D. B. B. Development of cardiac parasympathetic neurons, glial cells, and regional cholinergic innervation of the mouse heart. Neuroscience 221, 28–36 (2012).

19. McDavid, A. et al. Data exploration, quality control and testing in single-cell qPCR-based gene expression experiments. Bioinformatics 29, 461–467 (2013).

20. Hamilton, T. G., Klinghoffer, R. A., Corrin, P. D. & Soriano, P. Evolutionary divergence of platelet-derived growth factor alpha receptor signaling mechanisms. Mol. Cell. Biol. 23, 4013–25 (2003).

21. Hughes, E. G., Kang, S. H., Fukaya, M. & Bergles, D. E. Oligodendrocyte progenitors balance growth with self-repulsion to achieve homeostasis in the adult brain. Nat. Neurosci. 16, 668–76 (2013).

22. Ramilowski, J. A. et al. A draft network of ligand-receptor-mediated multicellular signalling in human. Nat. Commun. 6, 7866 (2015).

23. Bach, L. A. Endothelial cells and the IGF system. J. Mol. Endocrinol. 54, R1–13 (2015).

24. Bikfalvi, A., Klein, S., Pintucci, G. & Rifkin, D. B. Biological roles of fibroblast growth factor-2. Endocr. Rev. 18, 26–45 (1997).

25. Glebova, N. O. Heterogeneous Requirement of NGF for Sympathetic Target Innervation In Vivo. J. Neurosci. 24, 743–751 (2004).

26. Lohela, M., Bry, M., Tammela, T. & Alitalo, K. VEGFs and receptors involved in angiogenesis versus lymphangiogenesis. Current Opinion in Cell Biology 21, 154–165 (Elsevier Current Trends, 2009).

27. Clausen, B. E., Burkhardt, C., Reith, W., Renkawitz, R. & Förster, I. Conditional gene targeting in macrophages and granulocytes using LysMcre mice. Transgenic Res. 8, 265–277 (1999).

28. Madisen, L. et al. A robust and high-throughput Cre reporting and characterization system for the whole mouse brain. Nat. Neurosci. 13, 133–140 (2010).

29. Sudo, T. et al. Functional hierarchy of c-kit and c-fms in intramarrow production of CFU-M. Oncogene 11, 2469–76 (1995).

30. Conway, J. G. et al. Inhibition of colony-stimulating-factor-1 signaling in vivo with the orally bioavailable cFMS kinase inhibitor GW2580. Proc. Natl. Acad. Sci. 102, 16078–16083 (2005).

31. Nam, J. et al. Coronary veins determine the pattern of sympathetic innervation in the developing heart. Development 140, (2013).

32. Blenck, C. L., Harvey, P. A., Reckelhoff, J. F. & Leinwand, L. A. The importance of biological sex and estrogen in rodent models of cardiovascular health and disease. Circ. Res. 118, 1294–1312 (2016).

33. Fang, L. et al. Differences in inflammation, MMP activation and collagen damage account for gender difference in murine cardiac rupture following myocardial infarction. J. Mol. Cell. Cardiol. 43, 535–544 (2007).

34. Cavasin, M. A., Tao, Z.-Y., Yu, A.-L. & Yang, X.-P. Testosterone enhances early cardiac remodeling after myocardial infarction, causing rupture and degrading cardiac function. Am. J. Physiol. Heart Circ. Physiol. 290, H2043–50 (2006).

35. Langlais, D., Barreiro, L. B. & Gros, P. The macrophage IRF8/IRF1 regulome is required for protection against infections and is associated with chronic inflammation. J. Exp. Med. 213, (2016).

36. Sevilla, L. M. et al. Glucocorticoid receptor and Klf4 co-regulate anti-inflammatory genes in keratinocytes. Mol. Cell. Endocrinol. 412, 281–289 (2015).

37. Eddleston, J., Herschbach, J., Wagelie-Steffen, A. L., Christiansen, S. C. & Zuraw, B. L. The anti-inflammatory effect of glucocorticoids is mediated by glucocorticoid-induced leucine zipper in epithelial cells. J. Allergy Clin. Immunol. 119, 115–122 (2007).

38. Ronchetti, S., Migliorati, G. & Riccardi, C. GILZ as a Mediator of the Anti-Inflammatory Effects of Glucocorticoids. Front. Endocrinol. (Lausanne). 6, 170 (2015).

39. Baban, B. et al. The role of GILZ in modulation of adaptive immunity in a murine model of myocardial infarction. Exp. Mol. Pathol. 102, 408–414 (2017).

40. Ruffell, D. et al. A CREB-C/EBPbeta cascade induces M2 macrophage-specific gene expression and promotes muscle injury repair. Proc. Natl. Acad. Sci. U. S. A. 106, 17475–17480 (2009).

